# Transcriptomic and metabolomic characterization of antibacterial activity of *Melastoma dodecandrum*

**DOI:** 10.1101/2023.04.10.536307

**Authors:** Wee Han Poh, Nur Syahirah Ruhazat, Lay Kien Yang, Devendra Shivhare, Peng Ken Lim, Yoganathan Kanagasundaram, Scott A. Rice, Marek Mutwil

## Abstract

Antibacterial resistance poses a significant global threat, necessitating the discovery of new therapeutic agents. Plants are a valuable source of secondary metabolites with demonstrated anticancer and antibacterial properties. In this study, we reveal that Melastoma dodecandrum exhibits both bacteriostatic and bactericidal effects against Pseudomonas aeruginosa and Staphylococcus aureus. Treatment with plant extracts results in membrane damage and a reduction in Pseudomonas swimming and swarming motility. A comparative analysis of bacterial transcriptomes exposed to Melastoma extracts and four distinct antibiotics indicates that the extracts trigger similar transcriptomic responses as triclosan, a fatty acid inhibitor. Activity-guided fractionation suggests that the antibacterial activity is not attributable to hydrolyzable tannins, but to unidentified minor compounds. Additionally, we identified 104 specialized metabolic pathways and demonstrated a high level of transcriptional coordination between these biosynthetic pathways and phytohormones, highlighting potential regulatory mechanisms in plant metabolism.

## Introduction

Antimicrobial resistance (AMR) has been identified by the World Health Organization (WHO) as one of the top ten global health threats affecting humanity (Antimicrobial resistance). This problem is exacerbated by the lack of new effective antibacterials within clinical pipelines (Antimicrobial resistance). Breakthroughs in genomics and bioinformatics in the 90s enabled the identification of essential bacterial genes and targets. Consequently, pharmaceutical companies have attempted to generate new leads via target-based rational drug design. However, this strategy had limited success, partly due to an incomplete understanding of resistance mechanisms and factors influencing drug membrane permeability (Payne et al., 2007; Lewis, 2020; Tommasi et al., 2015). Alternatively, bioprospecting, the examination of natural sources for active bioproducts, may be considered in the search for new antibacterial compounds. To this end, the plant kingdom represents a rich source of plant secondary metabolites, many of which are known to have antimicrobial properties and have been used in traditional medicine to treat bacterial infections (Kessler and Kalske, 2018; Singh et al., 2018; Lal et al., 2020; Frey and Meyers, 2010; Ríos and Recio, 2005).

While bioprospecting may enable the discovery of more target-specific and structurally complex active biological compounds, it comes with associated challenges (Cushnie et al., 2020). For example, there may be difficulties in isolating and identifying active compounds from biological extracts which may be further compounded if there are multiple active compounds that contribute synergistically to the therapeutic activity of the extract. Similarly, the subsequent elucidation of the mechanism of action of an active compound may also be a tedious process (Atanasov et al., 2021). Additionally, the further use and industrial production of plant metabolites may be hindered by a limited understanding of their biosynthetic pathways and that the active metabolites may only be produced under specific conditions in specific organs (Atanasov et al., 2021; Kumar et al., 2019). Multi-omics methods, consisting of a combination of transcriptomics, metabolomics, genomics and/or proteomics, have been utilized in the drug discovery process and have been recognized as a useful tool to identify new targets and the mechanisms of lead compounds (Goff et al., 2020). The multi-omics strategy may also be used to discover plant metabolic genes and pathways using co-expression analysis and metabolite-based genome-wide association studies (Zhan et al., 2022; Zhao and Rhee, 2022; Julca et al.; Lim et al., 2022; Usadel et al., 2009). Taken together, such multi-omics approaches may represent the means to address challenges associated with bioprospecting and bridge the knowledge gap.

Members of the Melastoma genus, such as *Melastoma* malabathricum, *Melastoma candidum D. Don*, and *Melastoma dodecandrum,* have long been used in areas such as Malaysia, Taiwan, and China as traditional medicine for the treatment of dysentery, wounds, high blood pressure, diabetes, and skin diseases, amongst others (Joffry et al., 2012; Wang et al., 2008). More recently, studies have been carried out to scientifically evaluate *M. malabathricum* and *M. candidum D. Don* for acute toxicity, and antibacterial, antioxidant, and immunomodulatory activities (Mamat et al., 2013; Alnajar et al., 2012; Wong et al., 2012; Che Omar et al., 2013; Zheng et al., 2021). Similarly, efforts have been made to identify the chemical constituents of *M. dodecandrum* and its associated activities (Huang et al., 2021; Xu et al., 2023; Tong et al., 2019; Wang et al., 2017; Yang et al., 2014). However, limited studies have been carried out on evaluating the antimicrobial activity of the plant and identifying associated active compounds or mechanisms, with only a recent paper reporting moderate antibacterial activity of its ethanolic extract against diarrheagenic bacterial pathogens following an activity screen of 32 plants (Kudera et al., 2021).

Through the screenings conducted in this study, we have identified *M. dodecandrum Lour*. as a plant with antimicrobial and bactericidal effects on *Pseudomonas aeruginosa*. Phenotypic studies on *P. aeruginosa* indicate that the plant’s active compounds likely result in membrane damage and affect bacteria motility. Additionally, a comparative transcriptomic analysis of *P. aeruginosa* suggests that out of four antibiotics that target various vulnerabilities of bacteria, the plant extract’s mode of action overlaps with triclosan. Subsequent evaluation of fractions of the plant extract and activity testing of purified compounds suggests that an unknown compound is responsible for the antibacterial activity. By collecting samples of different organs, and carrying out transcriptomics studies and co-expression analysis, we investigated the biosynthetic potential of *Melastoma* and showed that the plant likely expressed many types of phenylpropanoids, with high levels of expression of sulfur-containing compounds, phenylpropanoids, and terpenoids. Finally, we showed co-expression patterns between the specialized metabolic pathways and hormones, suggesting that the pathways are transcriptionally linked.

## Methods

### Bacterial strains and culture conditions

*P. aeruginosa* PAO1 (ATCC BAA-47) was maintained on Luria-Bertani agar containing 10 g/L NaCl (LB10) (644520, Difco). *S. aureus* 25923 was maintained on TSB agar (236920, Difco). Cultures were rou tinely cultured in 10 mL LB10 media (244620, Difco) at 37°C with 200 rpm shaking overnight.

### Metabolite extraction

The flowers, fruits, leaves, stems, and roots of *M. dodecandrum* were sampled from the NTU Community Herb Garden. Leaves used solely for bioactivity screening were transported in dry ice and lyophilized using a freeze dryer (Labconco). Plant organs used for both bioactivity and transcriptomic studies were transported in liquid nitrogen. Subsequently, plant material was ground in liquid nitrogen to a fine powder. One hundred mg of the powder was aliquoted into a screw cap tube. Three 3mm steel beads and 500 µL of 80% HPLC-MS grade methanol were then added. The sample was subsequently homogenized for two cycles at 2000 rpm (PowerLyzer 24, Qiagen). The steel beads were removed, and the samples were placed on a shaker for 10 mins at 1500 rpm at room temperature in the dark. The samples were centrifuged at 10 min, 12000 × g. The supernatant was transferred into a new 2 mL Eppendorf tube and kept on ice. Five hundred µL of 80% HPLC-MS grade methanol was added to the remaining pellet, and the shaking and centrifuge step was repeated twice until 1.5 mL of supernatant was collected. The supernatant was dried completely using a centrifugal vacuum concentrator (Centrivap, Labconco) and dissolved in 100 – 500 µL of DMSO before use.

### Antimicrobial screening and activity testing

Dried methanol extracts derived from 100 mg of plant material or different plant organs were dissolved in 500 µL DMSO to give a stock solution of 200 mg/mL plant material. Overnight bacterial cultures were subcultured 1:10 in 10 mL of LB10 broth until the bacteria reached the exponential phase. Exponential phase bacteria were then diluted in Mueller-Hinton broth (MHB) to a final concentration of 5 × 10^5^ CFU/mL, and 2 µL of the stock DMSO solution was added to 100 µL of MHB to a final concentration of 4 mg/mL starting plant material or 2% v/v DMSO. Evaluation of HPLC fractions was carried out at 2% v/v DMSO, with each fraction corresponding to ∼10 mg/mL mass equivalent of the crude methanol extract.

### Extraction, bioassay guide fractionation, and chemical characterization

Freeze-dried leaves of M. dodecandrum (11 g) were milled and extracted twice with 400 mL of methanol (MeOH). The combined MeOH extract was dried under reduced pressure to afford 3 g of dried MeOH extract. Crude MeOH extract (2 g) dissolved in 5 mL of MeOH was fractionated using a Sephadex LH20 open column (5 x 35 cm). Compounds were eluted sequentially with 1 L of MeOH, 50% aqueous acetone, and acetone to generate 13 fractions. Biological testing results showed activity in only one fraction (58K, 120 mg) which was eluted in the acetone eluent.

50 mg of active fraction 58K was reconstituted in 15% aqueous MeOH, sonicated, and centrifuged. The insoluble pellet was collected (labeled as PPT, 5 mg) while the supernatant was injected into a Waters XTerra Prep MS C18 Column (10 x 300 mm) on an Agilent 1260 Infinity II Preparative HPLC system. The mobile phase consisted of water (A) and acetonitrile (B), both with 0.1% formic acid. The gradient elution started at 15% B for 5 mins, 15-30% B in 20 mins, 30-60% B in 25 mins, held at 60% for 10 mins, then increased from 60-100% B in 2 mins and washed at 100% B for 10 mins. The flow rate was 20 mL/min and the detection UV wavelength was at 254 nm. 8 fractions were generated from the preparative HPLC. All HPLC fractions, PPT and parent fraction 58K were submitted for biological evaluation.

Purified fractions were analyzed on an Agilent UPLC1290 coupled with a Quadrupole Time-of-Flight (Q-TOF) system. Separation was carried out with a reversed-phase C18 column (2.1 x 50 mm) at 0.5 mL/min, using a 10 mins linear gradient with 0.1% formic acid in both solvent A (water) and solvent B (acetonitrile). The typical QTOF operating parameters were as follows: negative ionization mode; sheath gas nitrogen flow, 12 L/min at 275°C; drying gas nitrogen flow, 8 L/min at 250°C; nebulizer pressure, 30 psi; nozzle voltage, 1.5 kV; capillary voltage, 1.5 kV. Lock masses in negative ion mode: TFA anion at m/z 112.9856 and HP-0921 TFA adduct at m/z 1033.9881. 1H NMR spectra were acquired on a Bruker DRX-400 NMR spectrometer with a 5-mm BBI Cryoprobe.

### Determination of minimum inhibitory concentrations

Minimum inhibitory concentrations (MIC) were determined using broth microdilution methods as described in (Wiegand et al., 2008). Briefly, plant extracts or equivalent concentrations of DMSO were diluted in MHB to four times the desired final concentration. 50 µL of the samples were added to 50 µL of MHB media on the first column of a 96-well plate, then serially diluted two-fold down the row. Untreated media-only growth control wells were also included. Overnight cultures of PAO1 were subcultured in LB10 medium and further incubated at 37°C with 200 rpm shaking. Exponential phase cells were then diluted in MHB to 1 × 10^6^ CFU/mL. Fifty µL of the bacterial culture was added to each well to a final concentration of 5 × 10^5^ CFU/mL, with the final concentration of plant extract falling between 0 – 50 mg/mL (0 – 5% DMSO). Sterility control wells, consisting of only MHB media, and blank control wells, containing 0 – 50 mg/mL of plant extract and MHB media, were also included. The 96-well plate was incubated for 24 h at 37°C with OD readings taken at 600 nm every 10 mins using a microplate reader (Tecan M200). Readings from blank control wells were subtracted from wells with bacteria added to account for high background readings at high plant extract concentrations. MIC was similarly determined for 0 – 512 µg/mL of the antibiotics and antimicrobial compounds triclosan, rifampicin, ceftazidime, and gentamicin. MIC was defined as the first concentration at which no growth is observed after 24 h incubation.

### Time-kill kinetics assay

Overnight cultures of PAO1 were subcultured in LB10 medium and further incubated at 37°C with 200 rpm shaking. Exponential phase cells were diluted to 1 × 10^6^ CFU/mL in MHB media and treated with 0 – 2 × MIC of plant extract. Vehicle controls were treated with equivalent volumes of 0 – 5% v/v DMSO. At each timepoint, samples were collected for serial dilution in PBS and CFU enumeration. Five µL of each dilution was spot plated onto LB10 agar plates and incubated at room temperature overnight. The first spots containing between 3 – 30 colonies were counted. The assay was carried out independently twice with two technical replicates each time.

### Evaluation of membrane integrity

The effect of plant extract on membrane integrity was evaluated using the LIVE/DEAD stain (L7012, Thermo Fischer) and microtiter plate reader (Tecan) as described in the commercial protocol. Briefly, sub-cultured exponential phase bacterial cells were washed in 1 × PBS (PBS). 5 × 10^7^ CFU/mL of bacterial cells diluted in PBS were subsequently treated with a non-inhibitory concentration of 0 – 4 mg/mL of filtered plant extract or equi-volume of DMSO for 10 mins. Samples were then treated for a further 30 mins with LIVE/DEAD stain readings taken with excitation at 485 nm and emission at 535 (red) and 635 nm (green). Dead cells were prepared via incubation of bacteria at 65 °C for 30 mins. The percentage of live cells is determined using a standard curve generated through a mixture of live and dead cells and the calculation of the green/red fluorescence ratio as stated in the commercial protocol. When LIVE/DEAD stain is used in microscopy, 2 × 10^7^ cells were seeded into a well of a poly-L lysine pre-treated ibidi 8-well chamber slide (80826, ibidi). The cells were allowed to attach for 10 mins before each well was washed and treated for 10 mins with 0 - 16 mg/mL of plant extract or equivolume of 0 - 1.6% v/v DMSO. Following treatment, each well was washed again and stained with LIVE/DEAD stain as described in the commercial protocol. At least three images per sample were taken using an epifluorescence microscope (Axio Observer Z1, Carl Zeiss) using the FITC filter for imaging SYTO 9 fluorescence and AF 568 filter for imaging PI staining. Images of fluorescence staining were analyzed using Image J.

### Motility assays

Swimming and swarming assays with *P. aeruginosa* were carried out similarly to (Ha et al., 2014b, 2014a). Briefly, 0.3% w/v or 0.5% w/v of 1X M8 agar (1X M8 solution: 42.3 mM Na_2_HPO_4,_ 22 mM KH_2_PO_4,_ 8.56 mM NaCl) were prepared for use in swimming and swarming assay respectively. Plant extracts or equivalent volumes of DMSO were added to the well with the highest concentration of the extracts, then serially diluted 4-fold down the wells before the agar solidified to a final concentration of 0 – 4 mg/mL plant extract. The agar was dried under laminar flow for 45 mins and used immediately after drying. Overnight cultures of PAO1 were used for inoculation. The agar was inoculated either by dipping an Eppendorf pipette tip into the culture and then stabbing it midway of the depth of the agar (swimming assay) or by spotting 1 µL of the culture on the surface of the agar at the centre of the well (swarming assay). Following inoculation, the plates were incubated at 30 °C overnight and the images were taken 16 h after inoculation. Swimming and swarming areas were assessed using Image J by setting an appropriate scale and selecting the region of interest.

### RNA isolation and sequencing (*M. dedecandrum*)

Following sample collection and grinding into a fine powder (as described in ‘Metabolite extraction’), RNA was extracted from 100mg of a sample using Spectrum™ Total Plant RNA Kit (Sigma) Protocol A following manufacturer’s instructions. Three biological replicates were collected for each adult organ, while single replicates were collected for the young flowers. Quality control of all extracted RNA was carried out by Novogene (Singapore). Each sample was evaluated for its quantity, integrity, and purity using agarose gel electrophoresis, Nanodrop, and Agilent 2100 Bioanalyzer. Library construction was performed by Novogene where mRNA was enriched from total RNA with oligo(-dT) magnetic beads. The library was then quantified with Qubit and real-time PCR and sequenced using Illumina NovaSeq 6000, with paired-end sequencing of 150 base pairs (bp) per read and a sequencing depth of approximately 60-70 million reads.

### RNA extraction and sequencing (*P. aeruginosa*)

Overnight cultures of PAO1 were subcultured and grown to mid-log phase. Cells were adjusted to OD=0.4 and then treated with equivolume of 2X antibiotic dissolved in MHB medium to a final concentration of OD=0.2, MIC=1X (or 25 mg/mL plant extract and 128 µg/mL triclosan), and a final volume of 500 µL. The samples were incubated at 37 °C statically for 30 mins. Following treatment, 1 mL of RNA protect reagent (76506, Qiagen) was added to 500 µL of the sample. The sample was treated as described in the commercial kit and the dried pellet was stored at −80 °C. RNA was extracted using the RNeasy plus mini kit (Qiagen) as per the kit’s instructions. Quality control of all extracted RNA was carried out by Novogene (Singapore). Each sample was evaluated for its quantity, integrity, and purity using Nanodrop, Agilent 5400 Bioanalyzer, and agarose gel electrophoresis. cDNA library was constructed and sequenced by Novogene. Novogene performed library construction with no ribosomal RNA depletion. Library quantification and sequencing were then performed as described above.

### RNA sequencing analysis (*P. aeruginosa*)

Low-quality RNA-seq reads were removed, and the remaining reads were trimmed with fastp (v0.23.2)(Chen et al., 2018). Reads were then aligned to the PAO1 genome (NC_002516.2) and read counts were obtained using EDGE-pro (Magoc et al., 2013). Due to poor overall read alignment, one DMSO-treated sample (replicate 1) was discarded (data not shown). One-third of the reads from the remaining two DMSO-treated samples were removed and collated to form a new third replicate. To evaluate sample-level variability, read counts were transformed using variance-stabilizing transformation, followed by hierarchical clustering of samples using Euclidean distance as the distance metric (Love et al., 2014). Differentially expressed genes (DEGs) between each antibiotic-treated sample compared to the DMSO control were identified with read counts using DESeq2. Genes with |⳩⳩⳩_*2*_ fold change| ≥ 1 and Benjamini-Hochberg (BH) adjusted p-value < 0.01 were considered significant (Benjamini and Hochberg, 1995) for multiple-testing correction. Gene ontology (GO) enrichment analysis was performed on each list of significant DEGs against the PAO1 GO biological process data set by applying Fisher’s Exact test with BH correction (adjusted p-value < 0.01)(The Gene Ontology Consortium, 2021; Mi et al., 2019).

### Melastoma gene expression data preprocessing

Quality control and read trimming of raw RNA-seq data were performed with fastp (v0.23.2). To obtain normalized transcript abundances (transcripts per million; TPM) of the RNA-seq samples, the reads were pseudoaligned against the coding sequences of the *M.dodecandrum* genome (PRJCA005299 from the National Genomics Data Center)(Hao et al., 2022) and quantified using Kallisto (v0.46.1)(Bray et al., 2016). To evaluate sample-level variability, samples were clustered using Spearman’s correlation as the distance metric - ⳩_⳩, ⳩_ = *1* - |⳩_⳩,⳩_|, where ⳩ and ⳩ are vectors comprising TPM expression profiles for all transcripts of a pair of RNA-seq samples respectively and ρ is the Spearman correlation coefficient. The final gene expression matrix was derived after filtering out genes with an average TPM of 0 across all organs. Genes were translated into protein sequences using the transeq module (EMBOSS v6.6.0.0)(Rice et al., 2000).

### Enzyme Annotation and Pathway Prediction

Enzyme annotation and pathway prediction were performed following the pipeline described by (Hawkins et al., 2021). Briefly, protein sequences were annotated with E2P2 (v4), pathway membership was predicted using PathoLogic in PathwayTools (v26) with default settings, and semi-automated validation of the final pathways was performed by SAVI (v3.1). All pathways assigned to the manual-to-validate list by SAVI were retained except for PWY0-501 and PWY-5723, following recommended validation steps by SAVI. From SAVI’s list of pathways to add, PWY-5173 was discarded as it is a deprecated ID in the current version of MetaCyc used by PathwayTools.

### Gene Co-expression and Pathway Subcluster Identification

To determine if genes within an SM pathway are more co-expressed than by chance, Pearson’s correlation coefficient (PCC) for each gene-pair within a pathway was first calculated. To simulate random pathways, a list of pathway-labeled gene-pairs was generated. The pathway labels were then randomly shuffled amongst the gene-pairs. The PCC for each gene-pair was calculated, and the median PCC of each random pathway was extracted. This was repeated 100 times.

To identify subclusters of a pathway, genes within a pathway were hierarchically clustered based on their pattern of expression across samples using the dendrogram function from scipy with default parameters (where the distance threshold is 70% of the maximum dendrogram distance). A representative subcluster was only identified for a pathway if the subcluster consisted of genes that represented at least 75% of the annotated reactions in the pathway in the plant (degree of completeness). As some pathways have multiple genes catalyzing the same reaction, subclusters of differing sizes can have the same degree of completeness. In these cases, the largest subclusters will be chosen as representative subclusters. For pathways with no representative sub-cluster, the original pathway was the representative ‘sub-cluster’ in downstream analyses.

### Co-expression analysis of connected pathways

To investigate patterns of inter-pathway coexpression within and between SM classes, we generated a pathway coexpression network. First, gene-pairs of all SM genes were calculated using the ‘pearsonr’ function from the scipy package. Then, p-values were corrected with the BH method (Benjamini and Hochberg, 1995). Gene-pairs with PCC :2 0.6 and BH-adjusted p-value (two-tailed) < 0.05 were considered significant. Subsequently, the gene nodes within the gene co-expression network were labeled with their respective SM pathways (as determined by pathway prediction). Multiple pathway labels will be assigned to genes that are predicted to be involved in multiple SM pathways. Labels of "Superpathways", defined as pathways that combine individual pathways, will only be assigned to a gene if the gene is not already labeled with their respective subpathways.

Next, to determine if the intra-pathway coexpression was better than random, the network was then subjected to the following permutation analysis: for each pathway, the number of intra-pathway edges was counted. Pathway labels were then randomly shuffled and the number of intra-pathway edges was counted. The following empirical p-value was calculated for each pathway:

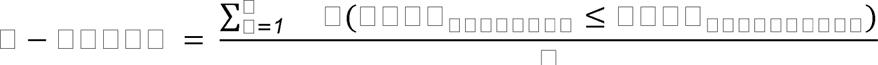

where *N* is the number of permutations, *I* is an indicator function that takes a value of 0 or 1 if the statement is false or true, respectively. This was repeated 10 000 times. Pathways with BH-adjusted p-values of < 0.05 were considered to be coexpressed (gene members of the particular pathway are coexpressed with each other). Inter-pathway coexpression between pathway-pairs were also determined similarly by counting inter-pathway edges that connect genes across any given two pathways.

To generate a pathway coexpression network, edges were used to connect SM pathways (represented as nodes) that were coexpressed with each other. Nodes were named according to the SM classes each pathway belongs to. Each node has a suffix of “_X” where X represents a single pathway from an SM class. e.g., if a PWY-A belongs to *Y* number of SM classes, then there will be *Y* number of PWY-A nodes in the network (“PWY- A_1, “PWY-A_2”, …, “PWY-A_*Y*”) while each edge joins 2 significantly coexpressed pathways.

## Results

### *Melastoma dodecandrum Lour.* shows antimicrobial activities

To assess the antimicrobial activity of *M. dodecandrum*, we sampled roots, petioles, leaves, vegetative branches, stems, fruits, and flowers from the Nanyang Technological University herb garden in Singapore (Figure 1A), and tested the antibacterial activity of methanol extracts derived from these organs. Bacterial growth inhibition studies indicated that the plant has antibacterial activity against gram-negative *P. aeruginosa* (Figure 1B, Table S1) and gram-positive *S. aureus* (Figure 1C, Table S2). When tested at 2% v/v (∼4 mg/mL extract) against *P. aeruginosa*, the stems showed the highest activity with a 33.13% (ANOVA, p < 0.0001) inhibition in growth over 18 h as compared to the DMSO controls, followed by flowers (32.56%), fruits (27.9%), leaves (15.54%), and roots (11.26%) (Figure 1B). Similarly, the stems showed the highest activity against *S. aureus* with 50.21% growth inhibition (p < 0.05)(Figure 1C). The vegetative branch, fruit, roots, and leaves showed moderate activity in a descending order ranging from 30.02 – 24.27% growth inhibition. In comparison, the flowers and petiole showed the lowest activity against *S. aureus* at 10.67 and 14.15% growth inhibition, respectively (Figure 1C).

**Figure 1.**
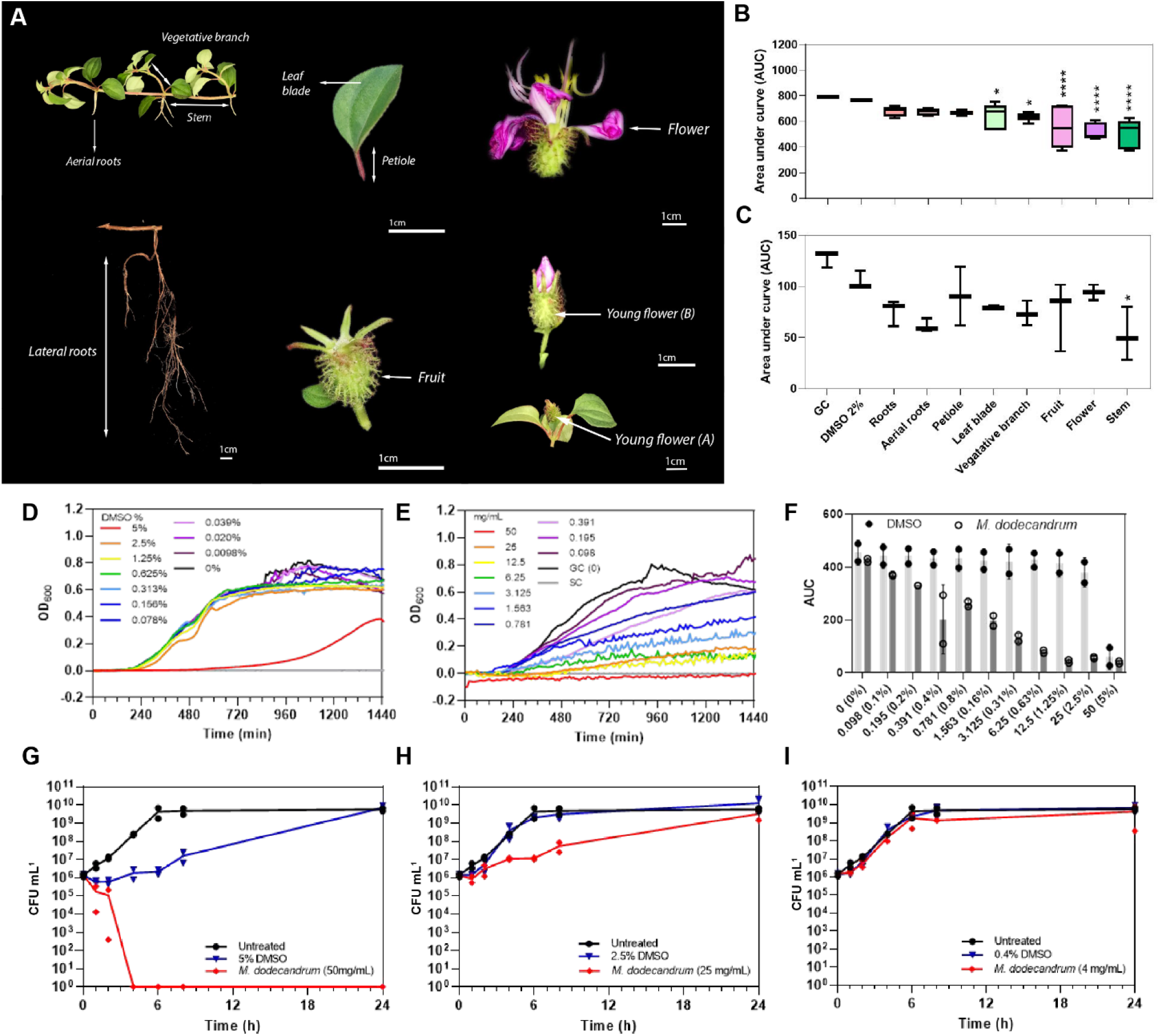
Anti-bacterial activities of *M. dodecandrum*. A) Various organs of M. dodecandrum Lour were collected for evaluating antimicrobial activity. The antibacterial activity of each plant organ against B) P. aeruginosa PAO1 and C) S. aureus 25923 was assessed using 2% v/v of plant methanol extract (∼4 mg/mL). The area under the growth curve (AUC) over 18 hours was measured and plotted. Boxplots represent data from at least three biological replicates, with differences in activity evaluated using one-way ANOVA with multiple comparisons against the 2% v/v DMSO control. D-F) Antibacterial activity of methanol plant extract from M. dodecandrum leaves was further evaluated at higher concentrations of 0-50 mg/mL (D) and equivalent DMSO concentrations of 0-5% v/v (E) over 24 hours. The AUC over 18 hours was plotted in (F), representing the mean from at least two biological replicates. G-I) The time-kill activity of 4-50 mg/mL of methanol plant extract from M. dodecandrum leaves or equivalent volumes of DMSO was assessed over 24 hours, with data points from each biological replicate plotted.

**Table 1:** Summary of the predicted enzymes and pathways.

| Total | Source/Tool | Number of genes/pathways (%) |
| --- | --- | --- |
| *Protein-coding genes* | Transcriptome | 32 021 |
| Predicted enzyme-coding genes | E2P2 | 8 035 (25.09) |
| Enzyme-coding genes in pathways |  | 3 867 (12.08) |
| Predicted SM genes | PathwayTools | 494 (12.77) |
| Predicted GM genes | PathwayTools | 3 097 (80.09) |
| Predicted S/GM genes |  | 276 (7.14) |
| *Pathways predicted* | PathwayTools, SAVI | 463 |
| Types of pathways |  | 9 |
| SM Pathways |  | 104 (3 have no genes assigned) |

*SM = Secondary Metabolite; GM = General Metabolite; S/GM = Genes that appear in both SM and GM pathways*

Next, we evaluated the minimum inhibitory concentration (MIC), defined as the concentration at which no growth is observed after overnight incubation. While leaves did not have the strongest activity amongst all organs (Figure 1B), they were the most abundant and easiest to collect. Hence, leaf methanol extract was used to evaluate the MIC of the extract on *P. aeruginosa* via a broth microdilution method. Complete growth inhibition was achieved when the extract was used at 50 mg/mL (Figure 1E – F, Table S3). The inhibitory effect was concentration dependent, with > 50% growth inhibition by 1.563 mg/mL of the extract compared to treatment with equivalent volumes of DMSO (Figure 1E-F, Table S4, Table S5). While 5% v/v DMSO treatment resulted in longer lag times, growth resumed by 720 min (Figure 1D).

The leaf methanol extract was bactericidal at 50 mg/mL, with complete cell death within four hours after treatment (Figure 1G, Table S6). In contrast, with the addition of equivalent 5% v/v DMSO, *P. aeruginosa* CFU recovered by 24 h (7.00 × 10^9^ CFU/mL) to levels comparable to that of untreated samples (5.65 × 10^9^ CFU/mL) (Figure 1G). A lower cell count was measured from 0 – 8 h with treatment with 25 mg/mL extract (Figure 1H), but the bacterial density reached untreated control levels by 24 h. Finally, *P. aeruginosa* treated with 4 mg/mL of the plant extract or equivalent concentrations of DMSO showed similar growth across all time points (Figure 1I), suggesting that a higher concentration of the plant extract is needed to achieve growth inhibition of bacteria.

### Melastoma extracts cause membrane damage and reduced swimming and swarming motility

To elucidate how the *M. dodecandrum* extracts affect the bacterial viability, we used LIVE/DEAD staining to assess potential membrane damage. The LIVE/DEAD stain consists of two nucleic acid staining dyes, propidium iodide (PI) and SYTO9. The green fluorescing SYTO9 can enter all cells, while the red fluorescing PI can only enter cells with compromised or damaged cytoplasmic membranes. PI, however, has a stronger affinity to DNA and hence can displace SYTO9 if present (Stiefel et al., 2015). While the kit is more accurately said to distinguish between cells with intact or compromised cell membranes, it is more often used to infer populations of living or dead cells under the assumption that all membrane-compromised cells are likely dead.

DMSO, when used at equivalent volumes of 0 – 0.8% v/v and a treatment time of 10 minutes, did not result in increased uptake of PI (Figure 2A, nearly all cells are green), and the proportion of live cells was calculated to be at ∼100% across all concentrations of DMSO (Figure 2B). In contrast, 10-minute treatment with 0 – 8 mg/mL of the plant extract resulted in a concentration-dependent increase in PI uptake and a corresponding reduction in the proportion of live cells from ∼100% in untreated samples to 13.6% in 8 mg/mL treated samples (Figure 2A, B, nearly all cells are red, Table S7).

**Figure 2.**
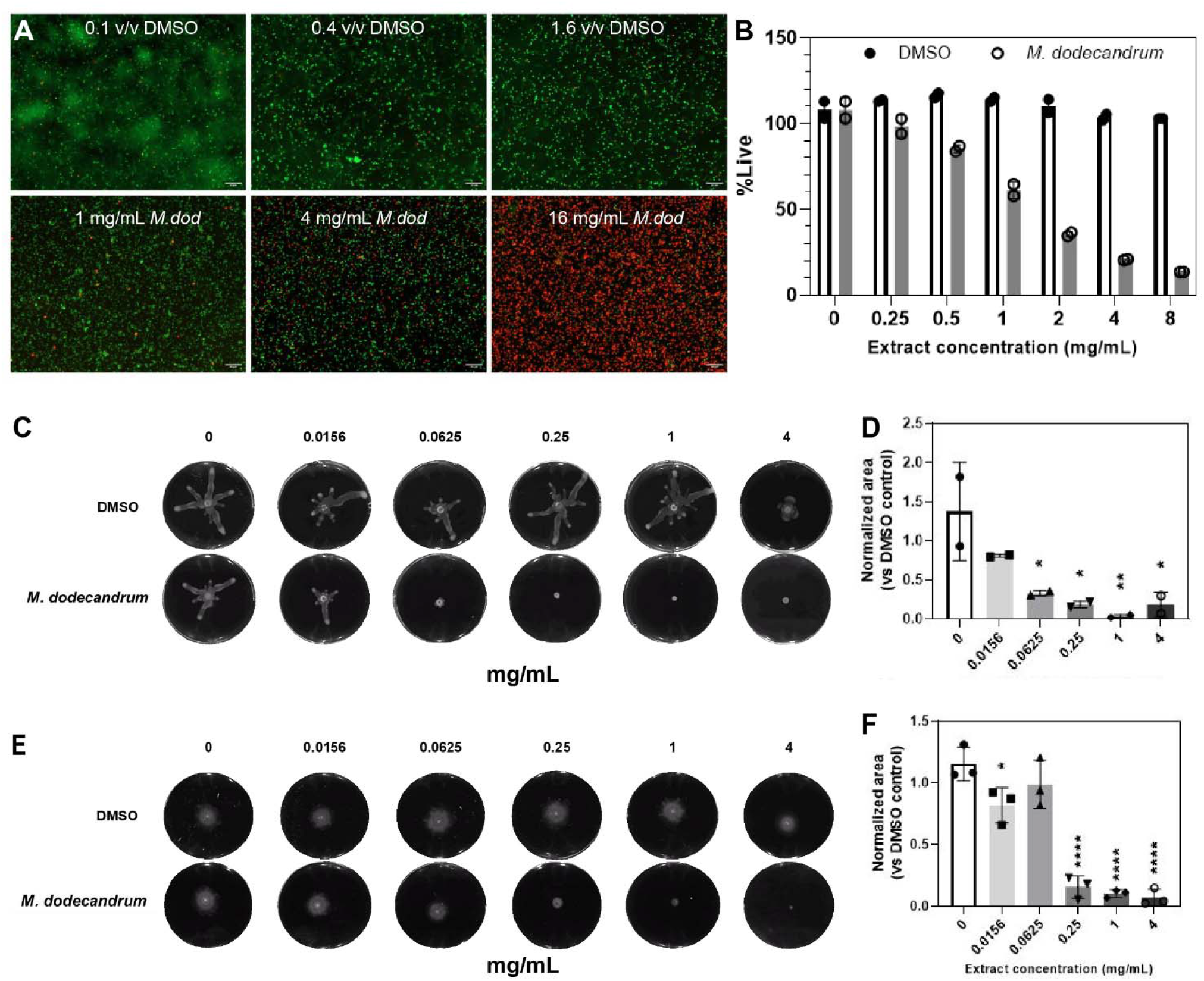
Viability, swimming and swarming analysis if Pseudomonas. A) Confocal microscopy images of P. aeruginosa PAO1 cells treated with equivolume of 0.1% (top left), 0.4% (top middle), or 1.6% v/v (top right) DMSO and 1 mg/mL (bottom left), 4 mg/mL (bottom middle), or 16 mg/mL (bottom right) M. dodecandrum extract and stained with LIVE/DEAD stain. Green and red cells indicate live and dead cells, respectively. B) Percentage of live cells following treatment with 0-8 mg/mL of M. dodecandrum leaf extract or an equal volume of DMSO, as determined using LIVE/DEAD stain and quantified using a microtiter plate reader. C) Swarming area experiment over 0-4 mg/mL M. dodecandrum extract ranges. D) Normalized area values from the swarming area experiments. E) Swimming area experiment over 0-4 mg/mL M. dodecandrum extract ranges. F) Normalized area values from the swimming area experiments. Each data point (Figure 2B, D, F) represents data from at least two independent experiments, with error bars representing the standard deviation of the mean. * = p _≤_ 0.05, ** = p _≤_ 0.01, **** = p _≤_ 0.0001.

Antibacterial compounds can also negatively affect the motility of bacteria, effectively disrupting their pathogenesis (Erhardt, 2016). Microscopy revealed reduced motility of cells treated with *M. dodecandrum* plant extract compared to the DMSO control groups, even at low concentrations of 1 mg/mL, where most *P. aeruginosa* cells appeared alive following 10 mins of treatment (Video 1 and 2 for DMSO control and extract treatment, respectively). To quantify and evaluate the effect of plant extract treatment on *P. aeruginosa* motility, swarming and swimming motility assays were carried out (Figure 2C – F). Swarming motility was significantly impaired at 0.0625 mg/mL (p < 0.05) (Figure 2C, D, Table S7), while swimming motility was impaired at 0.25 mg/mL (p < 0.0001) (Figure 2E, F, Table S7). These results indicated that *M. dodecandrum* leaf extracts exhibit a multi-pronged effect on *P. aeruginosa*, inhibiting viability, growth, and motility.

### Comparative transcriptomic analysis of antibiotic- and extract-induced gene expression changes in *P. aeruginosa*

Antibiotics can stimulate or depress gene expression in bacteria (Shitikov et al., 2022). Studying gene expression changes in responses to antibiotics can help the understanding of the mode of action of the various drug classes on bacterial adaptive ability, physiology, and metabolism. Since different antibiotics can elicit specific transcriptional responses (Hesketh et al., 2011), we hypothesized that gene expression analysis of *P. aeruginosa*- treated *M. dodecandrum* extracts and several antibiotics would allow us to propose the mode of action of the plant extracts.

To see if transcriptome changes caused by our plant extracts are similar to the changes caused by common antibiotics, we performed a comparative RNA sequencing analysis. We treated *P. aeruginosa* with rifampicin (RNA synthesis inhibitor), triclosan (fatty acid synthesis inhibitor), ceftazidime (cell wall synthesis inhibitor), gentamicin (protein synthesis inhibitor), *M. dodecandrum* extracts and DMSO control (Table S8).

Concentrations of extracts and antimicrobial compounds used for RNA-sequencing were selected based on results from microtiter plate growth curves (Figure S1). While 25 mg/mL of the extract did not fully inhibit *P. aeruginosa* growth by 24 h (Figure 1E, H), it was selected over 50 mg/mL as it inhibited growth to a similar extent within 18 h of growth without strong inhibition of growth associated with higher DMSO concentrations at 50 mg/mL (Figure 1F). Rifampicin, ciprofloxacin, and gentamicin were used at 1 × MIC of 32, 0.5, and 2 µg/mL respectively (Figure S1). Lastly, no growth inhibition was observed for triclosan over the range of concentration tested. Higher concentrations of triclosan treatment were not evaluated due to the precipitation of the compound in the media. As such, the highest concentration of 128 µg/mL, where some growth inhibition was observed, was used. DMSO was adjusted to the same concentration of 2.5% v/v for all samples to enable comparison with the control group. The resulting RNA-sequencing data was used to generate the gene expression matrix (Table S9).

Interestingly, the hierarchical clustering analysis of the 18 RNA-seq samples revealed that *M. dodecandrum* extracts show the highest Euclidean distance to all other samples (Figure 3A, *M. dodecandrum* RNA-seq samples form an outgroup). Similarly, rifampicin and triclosan also showed unique transcriptomic responses, while DMSO control, ceftazidime, and gentamicin displayed a similar response (Figure 3A). These responses could be explained by the number of significantly differentially up- (red) and down-regulated genes (Figure 3B, BH-adj. p-value < 0.01). *M.dodecandrum* elicited the highest number of up- (712) and down-regulated (757) genes, followed by rifampicin (331 up, 283 down) and triclosan (250, 135)(Table S10). Conversely, ceftazidime (40, 13) and gentamicin (99, 13) caused the mildest transcriptomic response. Overall, we observed that the Melastoma extracts showed some similarities of the differentially expressed genes with rifampicin and triclosan (Figure S2).

**Figure 3.**
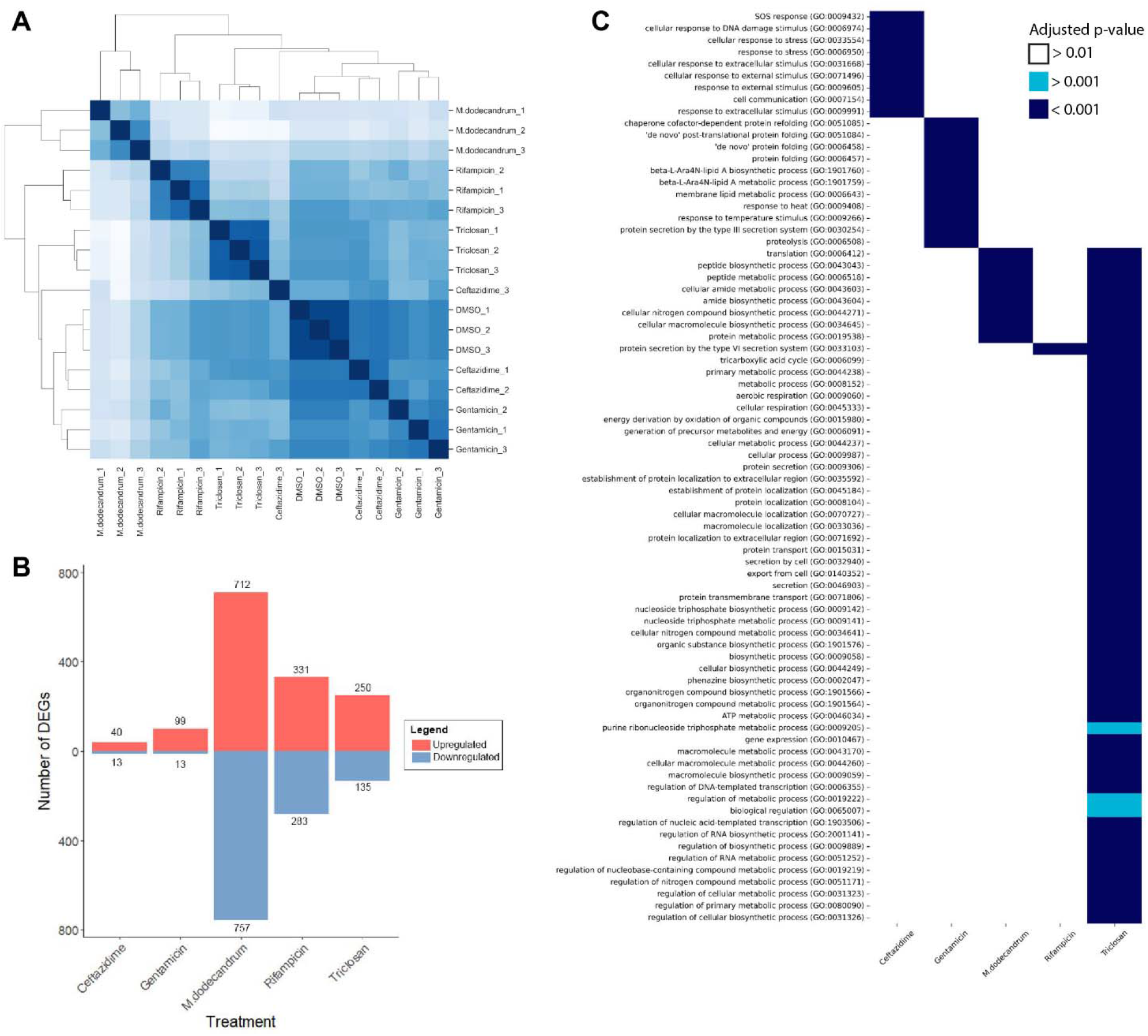
Gene expression responses of Pseudomonas aeruginosa to *M. dodecandrum* extracts and four antibiotics. A) Hierarchical clustering of RNA sequencing samples for DMSO, Melastoma extract, and four antibiotics. Each sample type is represented by three replicates, indicated by its suffix. B) The number of significantly differentially expressed genes (BH-adjusted p-value < 0.01), where up- and down-regulated genes are indicated in red and blue bars, respectively. C) Gene ontology enrichment analysis of the four antibiotics and Melastoma extracts. Antibiotics and the extract are shown in columns, while significantly enriched gene ontology terms are shown in columns. Cell colors indicate the different significance levels of the enrichment.

To see which biological processes are differentially expressed, we performed a Gene Ontology enrichment analysis (see methods). We observed that triclosan caused the highest number of differentially expressed GO terms (Figure 3C). These terms comprised many metabolic, biosynthetic, respiration, and macromolecule localization processes. The GO terms enriched in the plant extract-treated samples overlap with triclosan and are mostly metabolic and biosynthetic processes (translation, peptide, amide, protein, and nitrogen compounds). Conversely, rifampicin only induced changes in the type VI secretion system (Figure 5C). Interestingly, gentamicin caused changes in terms involved in the high-temperature response (folding, response to heat, and temperature stimulus), while ceftazidime responses were external stimulus and DNA damage (Figure 5C). Taken together, these results indicate the Melastoma extracts may have an overlapping mode of action with triclosan (fatty acid synthesis inhibitor).

**Figure 5.**
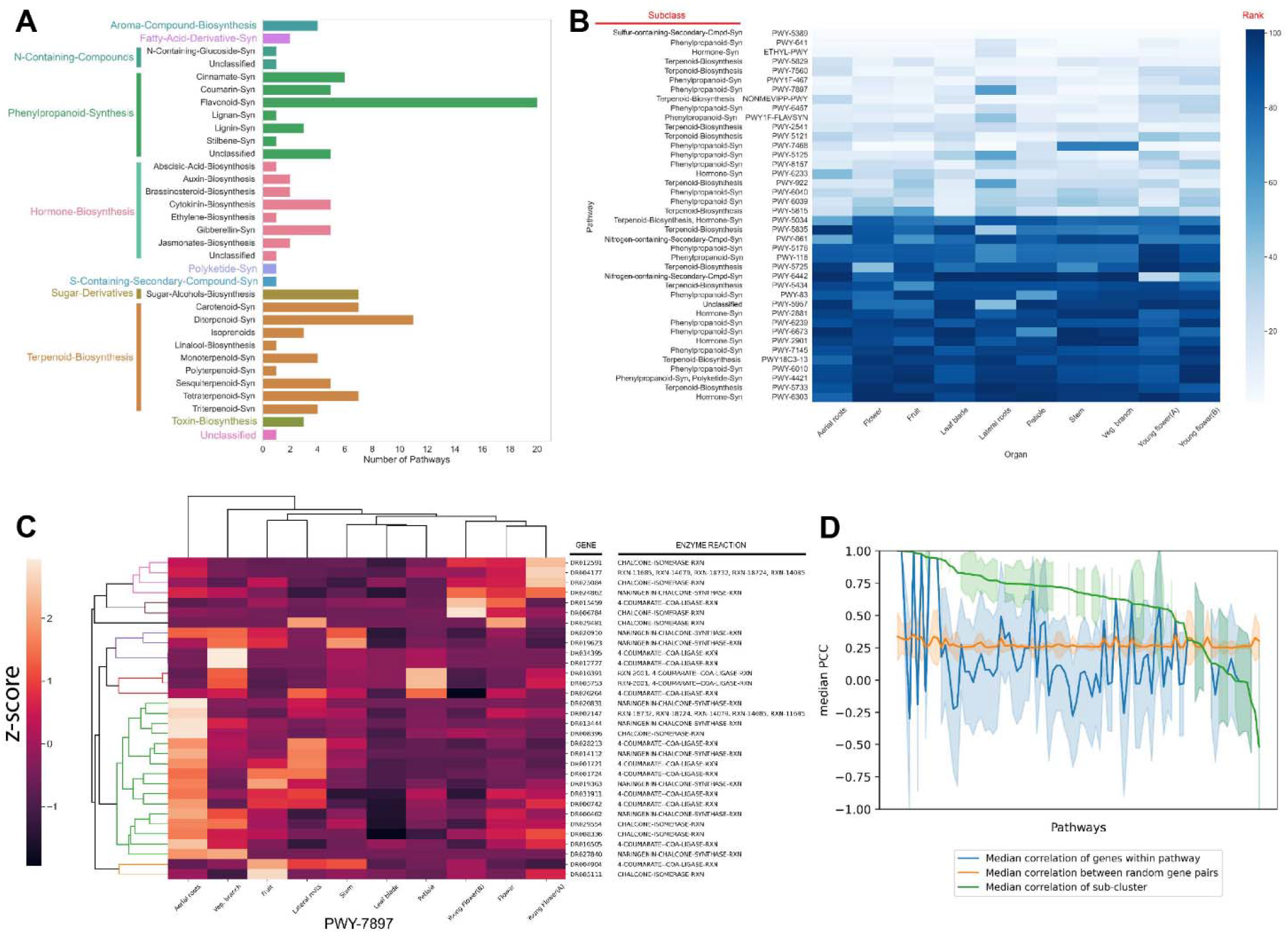
Identification of specialized metabolic pathways in *M. dodecandrum*. A) The number and types of putative specialized metabolic pathways identified by PathwayTools. B) Identification of the top 5 most and least expressed pathways from each organ. The scale represents the rank of a pathway (white = lowest rank, dark blue = highest rank). The rows represent pathways, while the columns indicate the sampled organs (as in Figure 1A). The lower rank (lighter color) indicates a higher expression. C) Clustering analysis of flavonoid biosynthesis pathway (PWY-7897). D) Correlation analysis of genes in the SM pathways. Pathways without confidence interval bands comprise two genes only.

### The antibacterial activity is conferred by unidentified metabolites

To identify the antibacterial compounds in Melastoma, we performed a bioassay-guided purification on the MeOH extract from the leaves. An active fraction (Parent) was obtained from Sephadex LH-20 column eluted at 100% acetone. It was subsequently redissolved in 15% aqueous methanol and centrifuged to separate the insoluble pellet (labeled as PPT, 5 mg). The supernatant was further fractionated by preparative reversed-phase HPLC and yielded 8 subfractions (A-H). Antimicrobial activity was observed in the Parent fraction and PPT, however, no activity was found in the A-H fractions (Figure 4A), indicating that the activity is caused by more than one synergistically-acting metabolites, or that the active metabolite was diluted during the fractionation.

**Figure 4.**
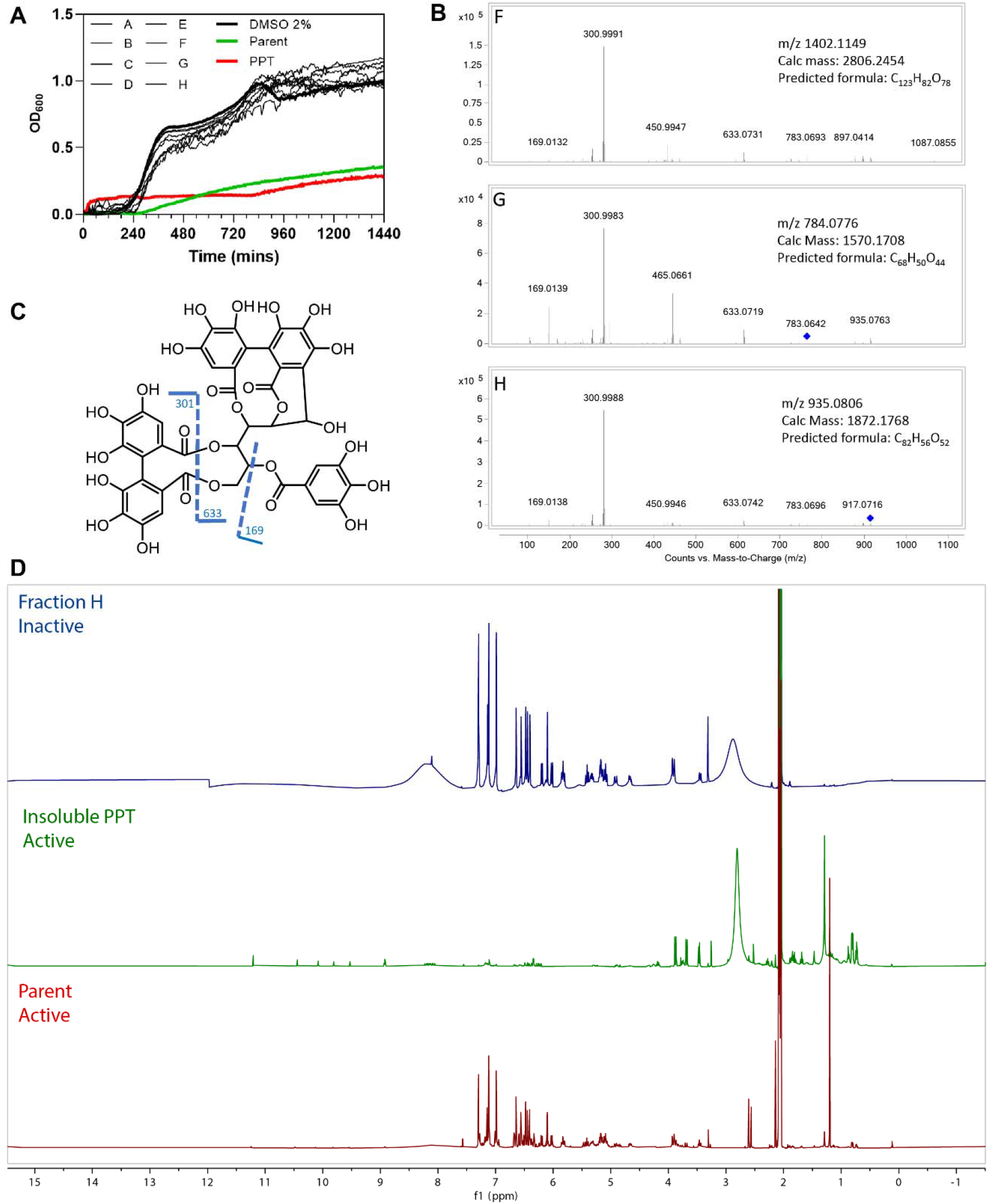
Bioassay results for HPLC fractions A-H, Parent fraction, and insoluble pellet (PPT). A) Antibacterial activity of eight HPLC fractions (A-H), Parent, insoluble pellet (PPT) and DMSO control. B) MS/MS profiles of the three fractions F, G, H. C) Fragmentation patterns of casuarinin. D) Overlay ^1^H NMR spectra of fractions H, PPT, and Parent, acquired in acetone-d_6_.

To study which metabolites might be present in the active fractions, we compared the active material (Parent and PPT) to the three inactive fractions F-H, using ^1^H NMR and LC-HRMS. HR-ESI-MS (High-resolution electrospray ionization mass spectrometry) of fractions F-H showed a major m/z of 1402.1149 [z=2]; 784.0776 [z=2] and 935.0793 [z=2], respectively (Figure 4B). A dominant fragment ion observed at m/z 301 indicated the existence of a hexahydroxydiphenoyl group (HHDP), which was characteristic of ellagitannins (Era et al., 2020). The MS/MS profiles were also in good agreement with that of hydrolyzable tannin casuarinin (m/z 935.0805 [z=1])(Figure 4C)(Chang et al., 2019). Coincidentally, fraction H presented the same m/z 935.0805, but a doubly charged [M-2H]^2-^. Therefore, the mass of H was calculated to be 1872.1738 and the molecular formula was determined as C_82_H_56_O_52_. From the predicted formula, fractions H, F (C_123_H_82_O_78_), and G (C_68_H_50_O_44_) were putatively identified as nobotanin analogs. The ^1^H NMR spectra (Figure 4D) of the major compound in fraction H also resembles spectra containing nobotanins (Chang et al., 2019). However, as none of the eight fractions A-H is active, it is unlikely that the tannins are significantly conferring antibacterial activity.

The overlay of ^1^H NMR spectra (Figure 4D) showed that the precipitation of the Parent fraction using 15% MeOH/H_2_O removed most of the hydrolysable tannins. ^1^H NMR of PPT revealed several unidentified signals in the Parent fraction, indicating a complex mixture consisting of phenolic compounds (9 - 11 ppm) and other unidentified compounds that could be responsible for the antibacterial activity.

### Gene expression analysis of specialized metabolic pathways in *Melastoma*

To investigate the types of metabolic pathways in Melastoma, we analyzed the 32021 protein-coding genes with PathwayTools (Karp et al., 2016). The analysis revealed that 25.09% (8035) of the genes are enzyme-coding and 12.08% (3867) could be assigned to a pathway (Table 1). Most of these genes were associated with general metabolism (GM, 3097 genes, 80.09% of all enzymes), followed by specialized metabolism (SM, 494 genes, 12.77%) and both GM/SM (276 genes, ∼7%, Table 1). From the 463 predicted pathways, 104 (22.4%) were associated with SM (Table 1, Table S11).

Next, we investigated the types of metabolites produced by the detected SM pathways. The majority of the compounds belonged to phenylpropanoids, of which flavonoids are most abundant (Figure 5A). Terpenoids, and more specifically diterpenoids biosynthesis comes in a close second. These observations are concurrent with previous studies which report these compounds in the larger Melastomacae family (Serna and Martínez, 2015).

To propose which specialized metabolites are most abundant in Melastoma, we analyzed the gene expression of the SM pathways. Since SM pathways are thought to be under strong transcriptional control (Mutwil, 2020), we set out to investigate the gene expression patterns in Melastoma. To this end, we isolated RNA from the same samples used to measure antibacterial activity (Figure 1A) and constructed a gene expression atlas for Melastoma (Figure S3, Table S12). We then ranked the SM pathways within each organ based on its average TPM expression. To identify the 20 highest and lowest-ranked pathways, we ordered them according to the sum of their ranks in each organ (Figure 5B). The top 5 pathways included a sulfur-containing compound, two terpenoids, a phenylpropanoid, and a hormone synthesis pathway. There are some pathways that show organ-specific expression (e.g. young-flower-specific nitrogen-containing secondary compound, PWY-6442).

Since SM pathways expand and diversify by gene duplications (Hofberger et al., 2013), the predicted SM pathways might likely represent several separate, related pathways biosynthesizing related metabolites. For example, several enzymes implicated in flavonoid biosynthesis (PWY-7897, Figure 5C) form an aerial root-specific cluster (green clade). This suggests that these genes biosynthesize a flavonoid in aerial roots, which might differ from flavonoids produced in flowers (first five enzymes, pink and brown clade, Figure 5C).

To investigate whether these functional sub-clusters are frequently found within the predicted pathways, we calculated the gene expression correlation (captured by Pearson Correlation Coefficient, PCC) within a pathway. We assumed that the median PCC should be high if the enzymes within a pathway are involved in the biosynthesis of the same metabolite. Surprisingly, while some of the pathways showed a high average PCC (blue line, Figure 5D), many pathways showed average PCC values like randomly chosen gene pairs (orange line, average PCC ∼ 0.25). Conversely, using gene expression data to identify the clusters containing >75% of annotated enzymatic reactions within a pathway resulted in a much higher average PCC value for nearly all pathways (green line, Figure 5D). This result shows that sequence similarity-based approaches such as PathwayTools should be supported by gene expression analyses for correct pathway inference.

### Transcriptional wiring of specialized metabolism in Melastoma

Melastoma might express over 100 SM pathways (Table 1) and metabolites (Figure 5A). Since these pathways should have specific functions, we expect the functionally-related pathways to be expressed at the same time and place. To identify such patterns, we investigate which pathways are more connected in the co-expression network than expected by chance. We first set a PCC threshold of ≥0.6, as the PCC values obtained from the randomized pathway assignments were all <0.6 (Figure 5D, randomized pathway assignment confidence interval is shown in orange). We then counted the number of co-expression edges connecting any two pathways and calculated which pathways were more connected than expected by chance (see methods). Interestingly, we observed high connectivity between the different pathways, where, e.g., most phenylpropanoid, terpenoid, and hormone biosynthesis pathways were significantly connected, indicating transcriptional coordination between the pathways (Figure 6A). Between 93 pathways, there were 251 significant connections.

**Figure 6.**
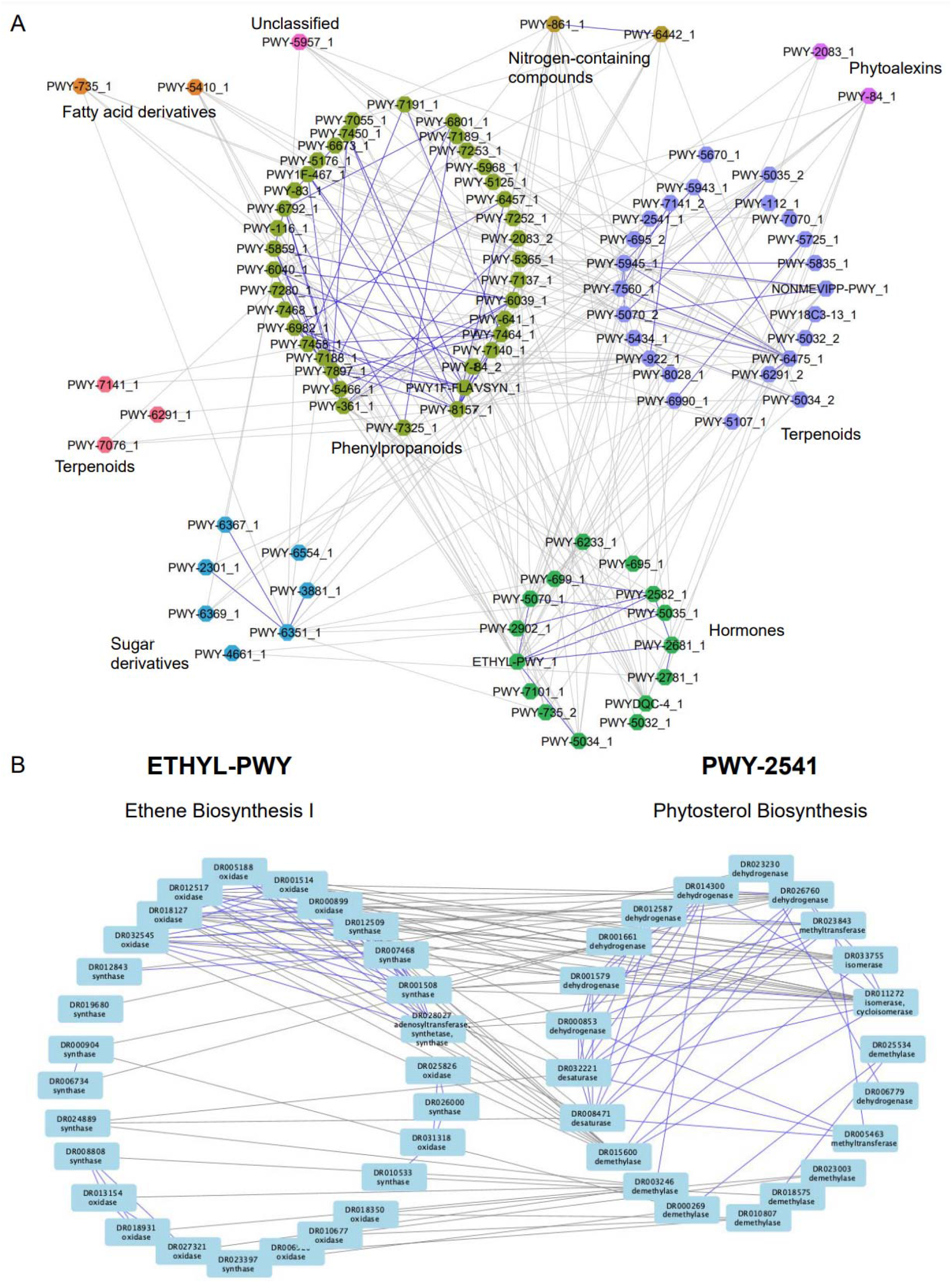
Co-expression network analysis reveals a transcriptional association between specialized metabolic pathways and hormones. A) Nodes represent pathways, while edges connect significantly connected pathways. Edges within the same pathway type are colored blue, while edges connecting different pathway types are colored gray. Node colors represent different pathway types. B) Ethylene (left) and phytosterol (right) biosynthesis pathways. Nodes represent enzymes, while edges connect enzymes correlated with r>0.6. Edges within the same pathway are blue, while edges connecting two pathways are gray.

To take a closer look at the connections that are found within and across the pathways, we investigated the gene co-expression networks of two connected pathways. The first pathway involves the biosynthesis of the plant hormone ethene (ethylene), which has numerous roles in plant development and stress responses (Chang, 2016). Ethene is synthesized from L-methionine by three enzymatic steps comprising methionine adenosyltransferase, 1-aminocyclopropane-1-carboxylate synthase (synthase), and 1- aminocyclopropane-1-carboxylate oxidase (oxidase) (https://pmn.plantcyc.org/PLANT/NEW-IMAGE?type=PATHWAY&object=ETHYL-PWY&detail-level=4). Interestingly, the co-expression network of the ethylene biosynthesis pathway shows several groups of co-expressed synthases and oxidases (Figure 6B, left pathway, blue edges connect co-expressed genes), suggesting that Melastoma contains several ethylene biosynthesis pathways. The other pathway represents the biosynthesis of phytosterol terpenoids, which comprise multiple steps that convert cycloartenol to stigmasterol, crinosterol, and brassicasterol in *Arabidopsis thaliana* (https://pmn.plantcyc.org/PLANT/NEW-IMAGE?type=PATHWAY&object=PWY-2541&detail-level=4). The Melastoma pathway contains several dehydrogenases, methyltransferases, isomerases, demethylases, and desaturases that modify the cycloartenol (or a related compound) to other sterols (Figure 6B, right pathway). While PWY-2541 has been characterized in Arabidopsis, it is likely that this pathway produces other, related compounds in Melastoma. Within each pathway, we also observed subclusters of genes that are co-expressed with the same gene(s) in the other pathway. While the analysis does not reveal any clear organ-specific clusters like in Figure 5C, this could suggest separate clusters of SM pathway expansion where subclustering occurs not only within a pathway but with other pathways too. The co-expression edges found between the ethylene and phytosterol pathways indicate that the two pathways are transcriptionally positively coordinated. This suggests that an increase in ethylene might result in increased biosynthesis of phytosterols. This is in line with a recent study that showed that exogenous ethylene led to a profound change in the ratio of stigmasterol to sitosterol (Markowski et al., 2022).

In conclusion, the high connectivity observed between various pathways, such as phenylpropanoid, terpenoid, and hormone biosynthesis pathways in Melastoma, highlights the intricate transcriptional coordination between these pathways, suggesting that they work together to regulate key aspects of plant development, stress responses, and secondary metabolite production, ultimately contributing to the plant’s adaptive capabilities and overall survival.

## Discussion

In this study, we showed that *M. dodecandrum* shows antibacterial activities against gram-positive and -negative bacteria (Figure 1). The plant is known to produce a diverse range of chemical compounds and metabolites, including flavonoids, tannins and ellagitannins, phenylpropanoids, long-chain fatty acids, aromatic acids, terpenoids, steroids, alkaloids, glycosides and monosaccharides (Zheng et al., 2021). Plant specialized metabolites can exert their antibacterial effects through various mechanisms. This includes but is not limited to, inhibition of bacterial nucleic acid and protein synthesis, inhibition of enzymes and respiration or metabolism, disruption of cell wall synthesis or cell membrane permeability or function, influencing the expression of virulence genes or chelation of essential metal ions such as iron (Payne et al., 2007; Tommasi et al., 2015; Lewis, 2020; Kessler and Kalske, 2018; Frey and Meyers, 2010; Singh et al., 2018; Ríos and Recio, 2005).

Results from LIVE/DEAD staining, in conjunction with viable CFU counts, suggest that the *M. dodecandrum* extract results in membrane damage (Figure 2). Specifically, following 10 min treatment with increasing concentrations of *M. dodecandrum* extract, LIVE/DEAD staining indicates an increasing proportion of dead cells at 16 mg/mL (Figure 2A, B). Conversely, viable cell counting suggests that there is no significant difference or drop in CFU compared to the control groups at 25 mg/mL (Figure 1H). Likewise, no significant differences in viable cell counts were observed in time-kill studies involving samples treated with 4 mg/mL w/v of the plant extract within an hour of treatment (Figure 1I). Recent studies suggest intermediate living/dead states in populations of bacteria stained with the LIVE/DEAD kit (Cushnie et al., 2020). Furthermore, there is evidence that, in some cases, PI staining may underestimate the viability of bacterial cells (Rosenberg et al., 2019). Taken together, it is likely that the increased PI staining following extract treatment is partly due to membrane damage and not actual cell death.

In addition to membrane damage, treatment with *M. dodecandrum* extract leads to reduced swimming and swarming motility in *P. aeruginosa* (Figure 2C-F, Video 1). Both forms of motility are flagella-mediated (Kazmierczak et al., 2015). The effects on motility may be related to the extract’s effects on the bacterial membranes, as flagella rotation is driven by ion flux and membrane-active antibiotics that result in the alteration of membrane and changes in cation permeability have been suggested to reduce flagella activity and in turn, motility in *P. fluorescens* (Manson et al., 1980; Faust and Doetsch, 1971). Similarly, cationic peptides that target cell membranes may also affect flagella integrity, and some cationic peptides can reduce swimming and swarming motility in *E. coli* (Faust and Doetsch, 1971). Unlike swimming motility, swarming motility is a much more complex and tightly regulated adaptation process involving the upregulation of virulence gene expression and antibiotic resistance (Overhage et al., 2008). In addition to functional flagella, swarming is also dependent on quorum sensing and type IV villi (Déziel et al., 2003; Köhler et al., 2000). Several plant metabolites have been reported to inhibit swarming via various mechanisms. For example, ginseng extract reduces swarming by reducing the production of QS signals in *P. aeruginosa,* while some flavonoids act as allosteric inhibitors of QS receptors (Wu et al., 2011; Paczkowski et al., 2017; Rütschlin and Böttcher, 2020). Some phenolic plant compounds or tannins can affect swarming but not swimming motility by affecting surfactant production (Rütschlin and Böttcher, 2020; O’May and Tufenkji, 2011). Beyond plant metabolites, antibiotics such as gentamicin and ciprofloxacin have also been reported to reduce swarming either via disruption of QS or through SOS responses that prevented polar chemosensory array formation (Rütschlin and Böttcher, 2020; Irazoki et al., 2017).

Comparison of gene expression responses of Pseudomonas treated with Melastoma extracts revealed that the plant extracts elicited the most unique and highest number of differentially expressed genes compared to four antibiotics (Figure 3A, B). This is not surprising, as the plant extract contains hundreds of different compounds that might synergistically affect different aspects of Pseudomonas biology. Surprisingly, the observed transcriptomics responses were not aligned with the primary mode of action of the antibiotics (Figure 3C). For example, while rifampicin inhibits RNA biosynthesis, the observed transcriptomic changes affect the unrelated type VI secretion system (Figure 3C). While these results indicate that transcriptomics might not be suitable to reveal the primary mode of action of antibiotics, the overlap of the differentially expressed biological processes suggests that Melastoma extracts might act similarly to triclosan (fatty acid biosynthesis inhibitor), which is in line with the observed membrane damage phenotypes (Figure 3C).

To identify the metabolite(s) responsible for the antibacterial activity, we performed activity-guided fractionation. However, none of the purified fractions showed activity (Figure 4A), suggesting that either the activity if caused by multiple metabolites that together cause the activity or that the active metabolite is poorly soluble. The latter is more likely, as the insoluble material showed antibacterial activity (Figure 4A). The ^1^H NMR analysis of the partially purified Parent fraction indicated the presence of hydrolysable tannins (Figure 4B), which are known to show broad anti-bacterial activities (Tomiyama et al., 2016), and other health-promoting activities due to their anti-oxidant properties (Abu Zarin et al., 2016). However, a deeper analysis using different fractionation methods showed that the activity was not due to the major hydrolysable tannins, but some other unidentified minor compounds. A larger scale of purification is needed to identify the active compounds.

The genomic and transcriptomic analysis revealed that M. dodecandrum contains many phenylpropanoid and terpenoid biosynthetic pathways (Figure 5A), and we were able to identify the most expressed pathways producing a sulfur-containing compound, two terpenoids, a phenylpropanoid (Figure 5B). Interestingly, within the predicted pathways, we observed several groups of highly co-expressed enzymes (Figure 5C, D), suggesting that these groups might represent several related pathways active in the different parts of the plant. Since enzymes biosynthesizing a given metabolite are expected to be expressed in the same cell, tissue and organ (Delli-Ponti et al., 2020), pathway inference approaches should be augmented with gene expression data. With >300,000 RNA-seq samples publicly available for hundreds of species (Julca et al., 2023), and several approaches that can be used to infer the biosynthetic pathways (Delli-Ponti et al., 2020), our and other groups can expect to provide more accurate inventories of specialized metabolism pathways.

Finally, we observed many co-expressed metabolic and hormonal pathways. This indicates that the specialized metabolic pathways are transcriptionally and functionally connected and that they are under hormonal control. This is in line with the existing knowledge linking many hormones to the biosynthesis of specialized metabolites (Markowski et al., 2022; Chen et al., 2019). Since co-expression analysis can identify hormone-pathway pairs (Figure 6A), our analyses can generate testable hypotheses of how plants’ hormones and metabolism are wired.

## Data availability

Raw RNA-seq data are deposited at the European Nucleotide Archive (ENA) under the accession number for bacteria (E-MTAB-12652) and plant (E-MTAB-12682).

## Supplemental Figures

**Figure S1.**
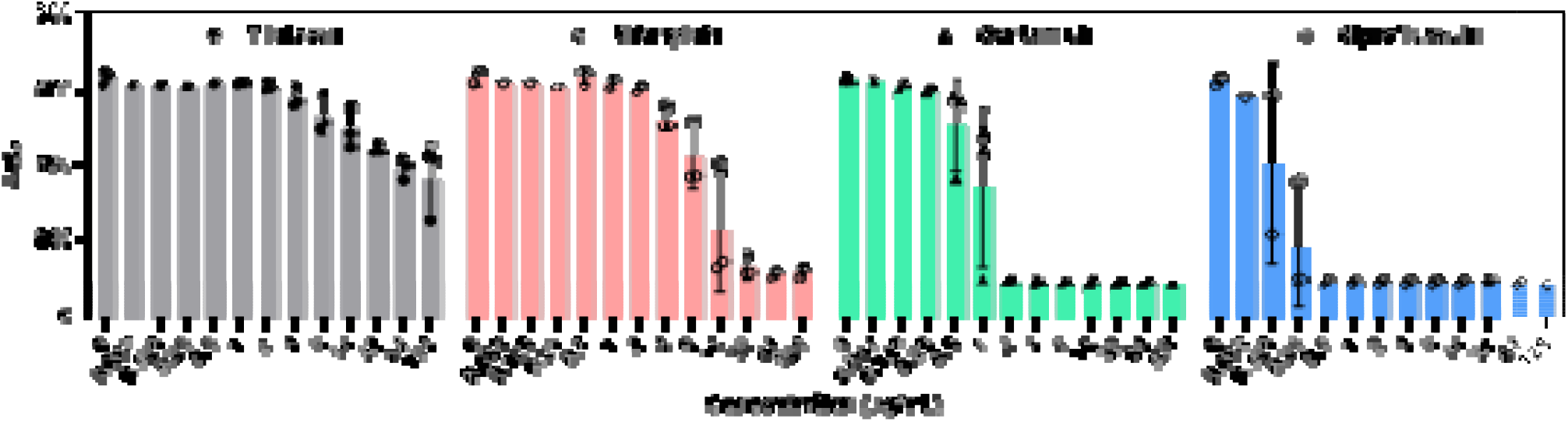
– Effects of various antimicrobials on the growth of *P. aeruginosa* PAO1. Effects of the treatment of *P. aeruginosa* with 0 – 128 µg/mL of the antimicrobials triclosan, rifampicin, gentamicin, and ciprofloxacin were evaluated and the area under the curve was plotted. Each data point represents data from one independent biological replicate while error bars represent the standard error of the mean. At least two biological replicates were carried out.

**Figure S2.**
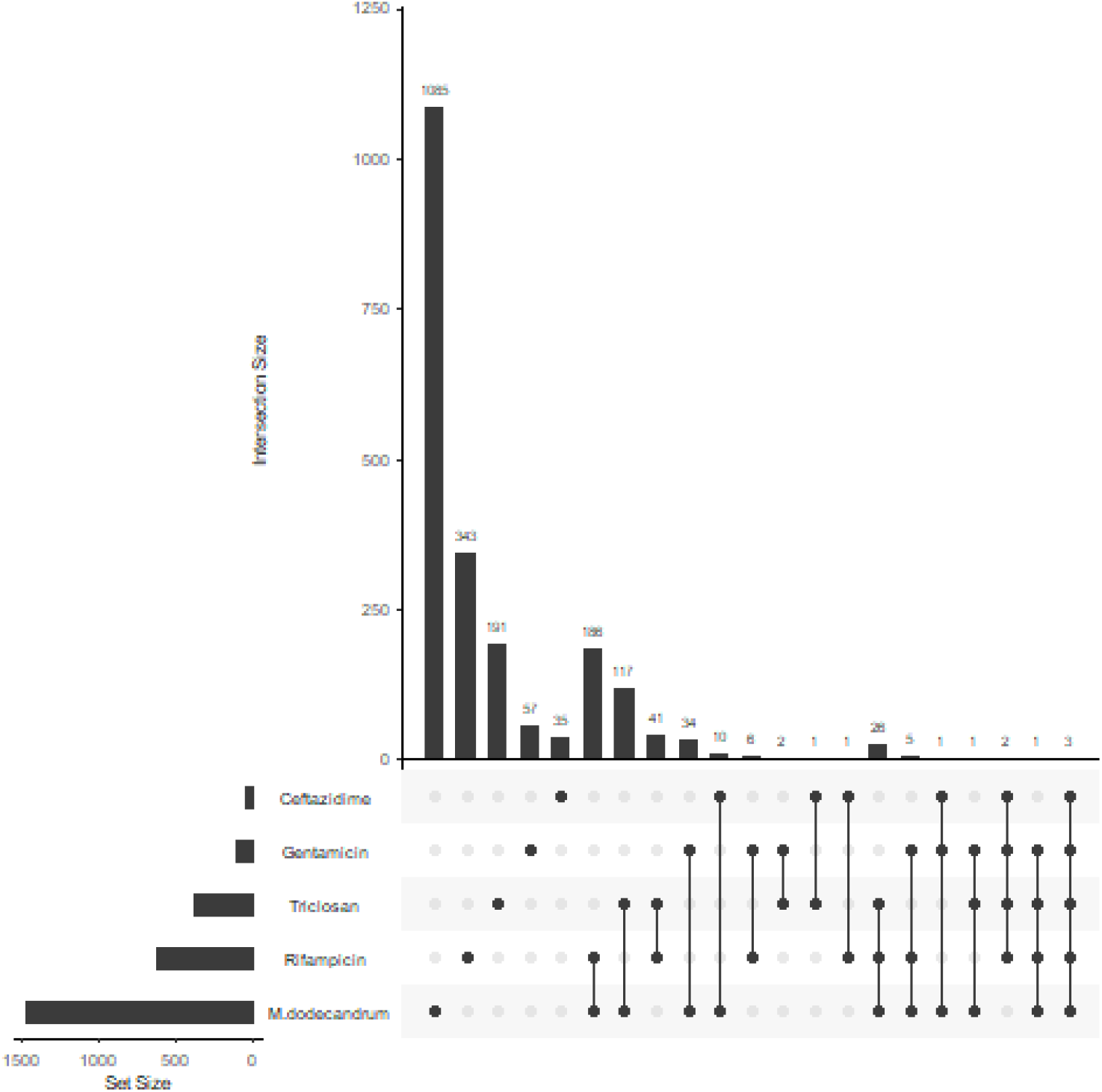
UpSet plot analysis of differentially expressed genes. The four antibiotics and *M. dodecandrum* plant extract is shown.

**Figure S3.**
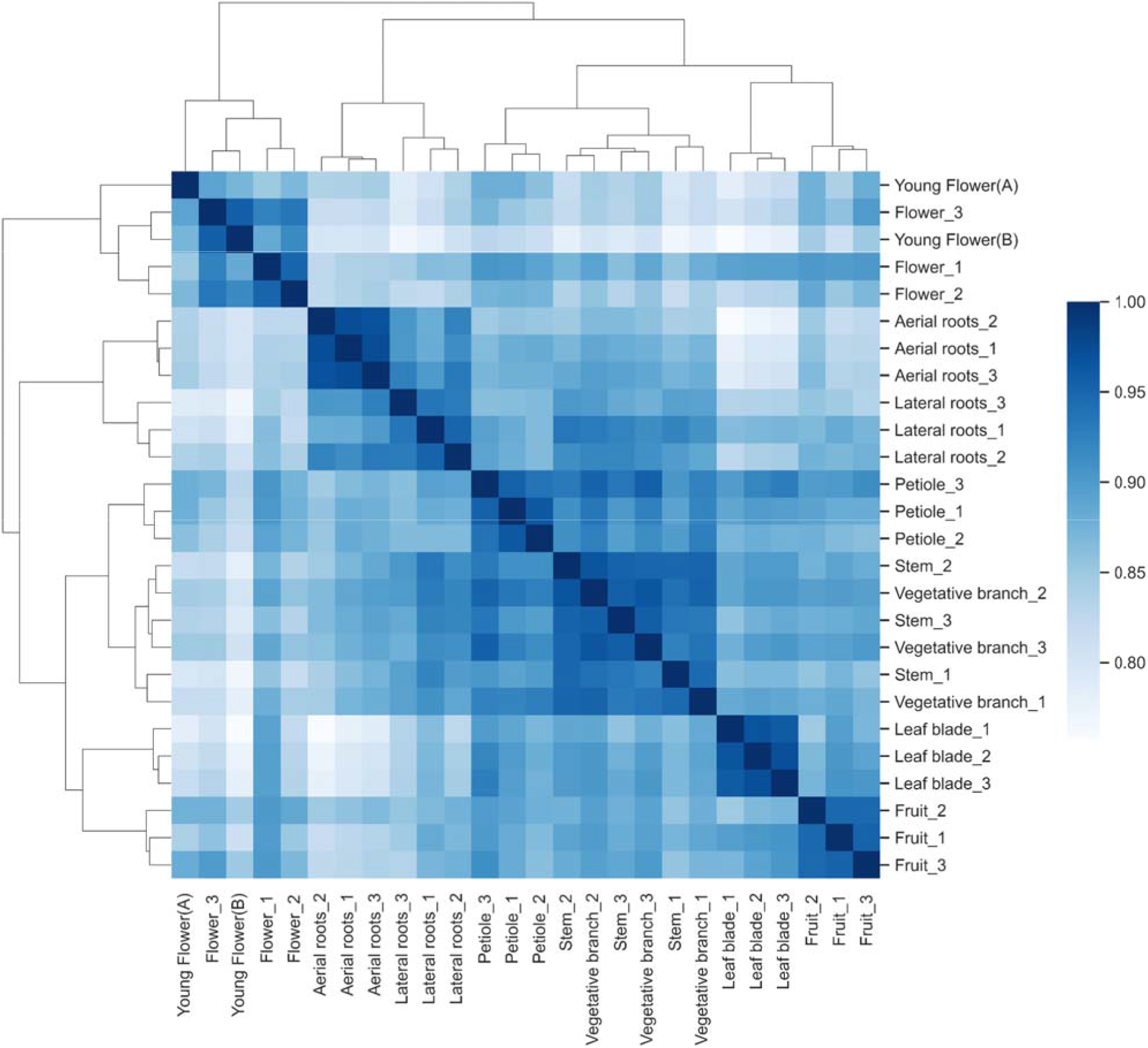
Hierarchical clustering analysis of gene expression data from Melastoma. Ward-linkage clustering based on Spearman correlation coefficient distance matrix.

## Supplemental Tables

**Table S1. OD600 values of Pseudomonas treated with the Melastoma extracts, control and DMSO.** The columns represent different organs, while rows represent timepoints.

**Table S2. OD600 values of Staphylococcus treated with the Melastoma extracts, control and DMSO.** The columns represent different organs, while rows represent timepoints.

**Table S3. AUC values at different concentrations of plant extracts (columns) and incubation times (rows).**

**Table S4. AUC values at different concentrations of DMSO (columns) and incubation times (rows).**

**Table S5. Summary of AUC values over different concentrations of DMSO and plant extracts.**

**Table S6. Time kill assay result over different concentrations of plant extracts and DMSO (columns) and hours of treatment (rows).**

**Table S7. Live/dead assay results.**

**Table S8. AUC values for different antibiotics (columns) and antibiotic concentrations (rows).**

**Table S9. Pseudomonas TPM expression matrix.** Rows represent genes, columns indicate samples.

**Table S10. Differential gene expression analysis result of Pseudomonas treated with the four antibiotics and plant extract.**

**Table S11. Putative pathways predicted by SAVI and PathwayTools, and their genes.**

**Table S12. Average expression of biosynthetic pathways across Melastoma organs.** Rows represent pathways, columns indicate organs, cells contain the average expression values in the organs.

## Supplemental Videos

**Supplemental Video 1. Pseudomonas treated with DMSO of equivalent volume to 1 mg/mL of plant extract.**

**Supplemental Video 2. Pseudomonas treated with 1 mg/mL of plant extract.**

## Author Contributions

W.H.P wrote the manuscript, performed bacterial wet- and dry-lab work with help from N.S.R, N.S.R. wrote the manuscript, performed plant wet- and dry-lab work with help from W.H.P., L.K.Y. did fractionation and structure elucidation of the active compound with help from Y.K, D.S. helped identify Melastoma, P.K.L. helped with bioinformatics analyses, S.R. supervised W.H.P., M.M. conceived the project, supervised W.H.P and N.S.R and helped with writing.

## Acknowledgments

We would like to thank NTU herb garden caretakers for providing the herb samples. The authors would like to thank the Mutwil Lab for useful discussions and suggestions during the lab meetings. Finally, we would like to thank Ing Tsyr Beh for help with the initial herb sample collection and screening.

